# Identification of CaVβ1 isoforms required for neuromuscular junction formation and maintenance

**DOI:** 10.1101/2025.06.10.658789

**Authors:** Vergnol Amélie, Bourguiba Aly, Bauché Stephanie, Traoré Massiré, Gelin Maxime, Gentil Christel, Pezet Sonia, Saillard Lucile, Meunier Pierre, Lemaitre Mégane, Perronnet Julianne, Tores Frederic, Gautier Candice, Guesmia Zoheir, Allemand Eric, Batsché Eric, Pietri-Rouxel France, Falcone Sestina

**Author notes:** These authors contributed equally to this work (Bourguiba Aly, Bauché Stephanie, Pietri-Rouxel France, Falcone Sestina).

## Abstract

Voltage-gated Ca²⁺ channels (VGCCs) are regulated by four CaVβ subunits (CaVβ1–CaVβ4), each showing specific expression patterns in excitable cells. While primarily known for regulating VGCC function, CaVβ proteins also have channel-independent roles, including gene expression modulation. Among these, CaVβ1 is expressed in skeletal muscle as multiple isoforms. The adult isoform, CaVβ1D, localizes at the triad and modulates CaV1 activity during Excitation-Contraction Coupling (ECC). In this study, we investigated the lesser-known embryonic/perinatal CaVβ1 isoforms and their roles in neuromuscular junction (NMJ) formation, maturation, and maintenance. We found that CaVβ1 isoform expression is developmentally regulated through differential promoter activation. Specifically, CaVβ1A is expressed in embryonic muscle and reactivated in denervated adult muscle, alongside the known CaVβ1E isoform. Nerve injury in adult muscle triggers a shift in promoter usage, resulting in re-expression of embryonic/perinatal Cacnb1A and Cacnb1E transcripts. Functional analyses using aneural agrin-induced AChR clustering on primary myotubes demonstrated that these isoforms contribute to NMJ formation. Additionally, their expression during early postnatal development is essential for NMJ maturation and long-term maintenance. These findings reveal previously unrecognized roles of CaVβ1 isoforms beyond VGCC regulation, highlighting their significance in neuromuscular system development and homeostasis.

## INTRODUCTION

Ca_V_β1 proteins, encoded by *Cacnb1* gene from exon1 to exon14, exist as several variants in multiple tissues. In skeletal muscle, *Cacnb1D* transcript, which spans exon1 to exon13end, gives Ca_V_β1D, the constitutive adult isoform. *Cacnb1E* transcript, ranging from exon2B to exon14, encodes Ca_V_β1E, an embryonic isoform that is re-expressed in adult skeletal muscle after nerve damage (1). Ca_V_β1 proteins have been originally described as key modulators of the skeletal muscle voltage-gated calcium (Ca^2+^) channel (VGCC) Ca_V_1 (2) and showed to be required for a proper Excitation Contraction Coupling (ECC). In particular, Ca_V_β1D, which localizes at the triads as subunit of Ca_V_1.1, likely holds the regulatory function for this process in adult skeletal muscle, modulating the interaction between the two Ca^2+^channels Ca_V_1.1 and Ryanodine Receptor 1 (RyR1) (1). In addition, Ca_V_β1 proteins have been reported as modulators of gene expression independently of Ca_V_1 channel, notably in muscle precursor cells regulating myogenesis, and in adult skeletal muscle after nerve damage, contributing to limit muscle mass loss (1, 3, 4). These roles are presumably accomplished by the embryonic Ca_V_β1E isoform, displaying nuclear localization (1, 3, 4).

In addition to the putative roles of Ca_V_β1D in regulating ECC at the triad and of Ca_V_β1E in modulating gene expression in the nucleus, another study highlighted the relevance of Ca_V_β1 proteins in skeletal muscle, by showing their crucial implication in the nerve-muscle connection during embryogenesis (5). During mammalian development and into early post-natal life, the formation, maturation and stabilization of neuromuscular junction (NMJ) are complex processes that require multiple cues for the establishment of functional nerve-muscle synapses. At early embryogenic stages, before the initiation of muscle innervation, and intrinsic to the muscle, both co-receptors LRP4 (low-density lipoprotein receptor-related protein 4) and MuSK (muscle-specific kinase) cluster together with clusters of aneural acetylcholine receptor (AChR) aggregates in a medial region of myotubes (6, 7). This phenomenon, known as muscle pre-patterning, represents the first initiating event of the NMJ formation (8). Alterations of aneural AChR pre-patterning give rise to abnormal growth and spreading of innervating neurites along the muscle fibers (5). Between E14 and E18, innervation by the motor neuron terminal drives the release of neural agrin, triggering the aggregation of AChR beneath the nerve terminal via Agrin/LRP4/ MuSK signaling. Later, depolarization induces the release of acetylcholine (ACh) from the nerve ending, which activates the postsynaptic AChR essential for NMJ maturation. In addition to being fundamental for NMJ formation and maturation, agrin/LRP4/MuSK signaling is also critical for NMJ maintenance and function throughout life. Moreover, Docking protein 7 (Dok7) one of the key players in maintaining NMJ integrity is indispensable for MuSK basal activity and activation of the tyrosine kinase domain by agrin (8, 9).

Ca^2+^ influx and release through Ca_V_1.1 and RyR1, rather than the ECC mechanism itself, are crucial for AChR aggregation during NMJ formation. This was demonstrated by using dysgenic (Ca_V_1.1-null) mouse embryos, which display abnormal AChRs distribution (10), which lacks both Ca^2+^ influx from Ca_V_1.1 and release from RyR1, show an abnormal AChR distribution (11). Similarly to these models, *Cacnb1^-/-^* mice show severely affected NMJ pre-patterning, independently of ECC (5). However, the role of Ca_V_β1 proteins in modulating various other aspects and mechanisms of NMJ formation, maturation and maintenance may have been underestimated.

In the present study, we aimed at identifying the major representative *Cacnb1* transcripts in embryonic pre- and post-innervated muscles in order to characterize their function in the formation and maturation of NMJ, as well as for NMJ maintenance implication in adult innervated muscles. We confirmed the expression of *Cacnb1D* in adult innervated muscle and found that *Cacnb1A*, in addition to *Cacnb1E*, is expressed in embryonic and denervated adult mouse skeletal muscle. We demonstrated that the expression of the different *Cacnb1* isoforms is driven by the activation of specific promoters on embryonic and adult muscles, and that denervation is associated with the restoration of the embryonic epigenetic program. Analysis of the function of embryonic isoforms in the formation and maturation of NMJs after ablation of the embryonic Ca_V_β1 isoforms revealed a precocious differentiation of primary myotubes *in vitro*, associated with an increase in the size of neural agrin-triggered AChR aggregates, while it induced a decrease in the size of AchR areas in early postnatal myofibers. In adult innervated muscle, Ca_V_β1 embryonic isoforms ablation led to NMJ destabilization, accompanied by denervation-like signs, accordingly to what observed in the early phases of muscle aging (12).

Overall, our findings reveal the crucial relevance of Ca_V_β1 embryonic isoforms in processes needed for the establishment, maturation and stability of functional NMJ, shedding light on novel fundamental players involved in development, growth and maintenance of the neuromuscular system.

## MATERIAL AND METHODS

### Plasmids & Adeno Associated Virus (AAV) production

pSMD2-shEx2A was already described (1) to specifically targets *Cacnb1* exon 2A (target sequence CCTCGGATACAACATCCAACA), therefore downregulating *Cacnb1* embryonic isoforms specifically. pSMD2-shEx2A has been prepared in AAV1 serotype vector by the AAV production facility of the Center of Research in Myology, by transfection in 293 cells as described previously (1). The final viral preparations were kept in phosphate-buffered saline (PBS) solution at −80°C. The particle titer (number of viral genomes) was determined by quantitative PCR. Transduction of this AAV1-containing plasmid will be referred shEx2A treatment.

### Animals and ethics

All animals were housed under specific-pathogen-free (SPF) conditions, with a 12 hours light cycle with access to rodent food and water ab libitum. Embryos were from breeding housed directly in UMS28 animal facility or from gestating females from Janvier Lab. Newborn (7 days-old) male and female mice were from breeding housed directly in UMS28 animal facility. Adult (2 months old at the beginning of the procedures) C57/BL6 female mice were from Janvier Lab and accustomed to the animal facility for one week before inducing the experimental procedures. In the end of the procedures, 7days old mice were euthanized by decapitation while adult mice were euthanized by cervical dislocation. Animal experimental procedures were performed in accordance with national and European guidelines for animal experimentation, with the approval of the institutional ethics committee (MENESR: Project #17975 and Project #45606).

### Injection procedures

For experiment on adult mice, 8 weeks-old female animals were injected with AAV1-shEx2A or AAV1-scramble in Tibialis Anterior (TA) muscles. Anaesthesia was achieved using isoflurane (3% induction, 2% maintenance), analgesia by buprenorphine (vetergesic 1mg/Kg, subcutaneous). One intramuscular injection (40µl/TA or 100µl/gastrocnemius (GAS) + Soleus (Sol)) was performed at a final titer of 1.10^9^ vector genomes (vg)/mg of muscle. Mice were sacrificed 8 weeks after the injection. For experiments on early stage muscle development, animals (female and male) were injected at 7 days old with AAV1-shEx2A or AAV1-scramble in TA and GAS/Sol muscles. Anaesthesia was achieved using isoflurane (4% induction, 4% maintenance). One intramuscular injection (10 μl/TA or 15µL/GAS+Sol) was performed at a final titer of 1.10^9^ vg/mg of muscle. Mice were sacrificed 3 weeks after the injection. Muscles were collected and either fixed in paraformaldehyde (PFA) 4% or frozen in isopentane precooled in liquid nitrogen and stored at −80 °C until histology or molecular analysis.

### Denervation procedures

Right sciatic nerves were neuroectomized (ablation of a 5-mm segment of the sciatic nerve) under general anesthesia (Isofluorane, 3% induction, 2% maintenance) and under buprenorphine (vetergesic 1mg/Kg, subcutaneous). Mice were sacrificed 2, 3 or 14 days after denervation and muscles were collected and either fixed in PFA 4% or frozen in isopentane precooled in liquid nitrogen and stored at −80 °C until histology or molecular analysis.

### Functional analyses

The force measurement of TA was evaluated by measuring *in vivo* muscle contraction in response to nerve stimulation, as previously described (1). Mice were anesthetized with Isofluorane, 3%, the knee and paw were fixed in place, and the distal tendon of the muscle was attached to the lever arm of a servomotor system (305B, Dual-Mode Lever, Aurora Scientific) using a silk ligature. Data were analyzed using the PowerLab system (4SP, ADInstruments) and software (Chart 4, ADInstruments). The sciatic nerve was stimulated using supramaximal square-wave pulses of 0.1ms in duration. Capacity for force generation was evaluated by measuring the absolute maximal force that was generated during isometric contractions in response to electrical stimulation (frequency of 75–150Hz; train of stimulation of 500ms). Maximal isometric force was determined at L0 (length at which maximal tension was obtained during the tetanus). Force was normalized by muscle mass as an estimate of specific force.

For electroneuromyogram (ENMG), two monopolar reference and recording electrodes were inserted, one close to the patella tendon and the other in the middle of the TA muscle. A monopolar ground electrode was also inserted into the contralateral hindlimb muscle. The sciatic nerve was stimulated with series of 10 stimuli at 5, 10, 20, and 40Hz. Compound muscle activity potential (CMAP) was amplified (BioAmp, ADInstruments), acquired with a sampling rate of 100kHz, and filtered with a 5kHz low-pass filter and a 1Hz high-pass filter (PowerLab 4/25, ADInstruments), and peak-to-peak amplitudes were analyzed with LabChart 8 software (ADInstruments).

During all experiments, mouse body temperature was maintained at 37°C with radiant heat.

### Primary cells, transduction and high differentiation induction

Primary myoblasts from WT newborn mice were prepared as described (13). Briefly, after hind limb muscle isolation, muscles were minced and digested for 1.5h in PBS containing 0.5mg/ml collagenase (Sigma) and 3.5 mg/ml dispase (GibcoLife Technologies). Cell suspension was filtered through a 40μm cell strainer and preplated in IMEM + 10% FBS + 0.1% gentamycin for 4h, to discard the majority of fibroblasts and contaminating cells. Non-adherent myogenic cells were collected and plated in IMDM + 20% FBS + 1% Chick Embryo Extract (MP Biomedical) + 0.1% gentamycin in 12-well plate or fluorodishes coated with IMDM 1:100 Matrigel Reduced Factor (BD Bioscience). Differentiation was triggered by medium switch in IMDM + 2% horse serum + 0.1% gentamycin. A thick layer of Matrigel Reduced Factor (1:3 in IMDM) was added 24h after differentiation medium addition. Myotubes were treated with 100µg/ml of agrin, and the medium was changed every 2 days, for 8 days of differentiation (13). Downregulation of embryonic Ca_V_β1 isoforms was achieved concurrently with the induction of differentiation through transduction of AAV1-shEx2A.

### RNA isolation and gene expression analyses

Total RNA was extracted from muscle cryosections using Trizol (ThermoFisher Scientific) /Direct-zol RNA MiniPrep w/ Zymo-Spin IIC Columns (Ozyme) and from cells using NucleoSpin RNA Columns (Macherey-Nagel). 200ng of total RNA were subjected to Reverse transcription (RT) using Superscript II Reverse transcriptase kit (18064022, ThermoFisher Scientific) or 1µg of total RNA using Maxima H Minus Reverse Transcriptase (EP0753, ThermoFisher Scientific), with a mix of random primers (48190011, ThermoFisher Scientific) and oligo(dT) (18418020, ThermoFisher Scientific), to generate complementary DNA (cDNA). 2µL of cDNA was amplified in 20µL reactions in PCR Master Mix (M7505, Promega), Phusion High-Fidelity PCR Master Mix with GC Buffer (F532L, ThermoFisher Scientific) or Q5 Hot Start High-Fidelity 2X Master Mix (M0494S, New England Biolabs) for RT-PCR and RT-PCR triplex. 2µL of 1:5 diluted cDNA was amplified in 10µL reactions in Power SYBR Green PCR MasterMix (ThermoFisher Scientific) to performed quantitative real-time PCR (qPCR) on StepOne Plus Real-Time PCR System (Applied Biosystems). Data were analyzed using ΔΔCT method and normalized to RPLPO (Ribosomal protein lateral stalk subunit P0) mRNA expression. The sample reference to calculate mRNA fold change is indicated in each panel. Primers are listed in **Table S1**.

### Nanopore sequencing

600ng of total RNA were subjected to RT with an anchored poly-dT primer (12577011, ThermoFisher Scientific) and Maxima H Minus Reverse Transcriptase. The amplification of the *Cacnb1* transcriptome was independently processed in each sample through two PCR steps, using 2μL of diluted RT reactions (12-fold dilution). First, a pre-amplification was performed for 20 cycles using specific primers fused with universal sequences U1 and U2 (referenced in **Table S2**). These primers are specifically base paired with the first and last exons of *Cacnb1* transcripts. The PCR reactions were treated with exonuclease I to remove primer excess and purified with the NucleoMag® NGS Clean-up and Size Select (Macherey-Nagel). Next, a second amplification of 18 cycles was performed to incorporate barcodes associated with individual samples. All samples were combined to create a stoichiometric multiplexed library, which was prepared using the Oxford Nanopore SQK-LSK109 kit. The library was subsequently sequenced using a MinION Flow Cell (R9.4.1). The Oxford Nanopore data were analyzed as previously described (14) and Fastq files generated in this study have been deposited in the ENA database (https://www.ebi.ac.uk/ena/browser) under accession code PRJEB89914.

### ChIP experiments

TA and GAS muscles were harvested from 8-weeks-old female mice, innervated or 2 days after denervation. Tissues were finely minced with a scalpel in PBS, incubated for 10min in PBS containing 1% formaldehyde, then 5min in PBS containing 125mM Glycine. Aliquots used for RNAPII immunoprecipitation were fixed a second time with Disuccinimidyl glutarate (DSG) diluted to 2mM in PBS for 40min at room temperature, as previously described (15). Tissues were then dissociated with a MACS dissociator using the muscle-specific program.

Resuspended material was incubated on ice 5min in 1mL of chilled buffer A (0.25% TRITON, 10 mM TRIS pH8, 10mM EDTA, 0.5mM EGTA), 30min in 1mL of buffer B (250mM NaCl, 50mM TRIS pH8, 1mM EDTA, 0.5mM EGTA), and then resuspended in 100 to 200µL of buffer C (1% SDS, 10mM TRIS pH8, 1mM EDTA, 0.5mM EGTA) at room temperature. All buffers were extemporaneously supplemented by Complete protease and PhosSTOP phosphatase inhibitors (Roche). Cell suspensions in 0.6mL µtube were sonicated in water bath at 4°C during 15min (15sec. ON, 15sec. OFF) with a Bioruptor apparatus (Diagenode) set on high power and then clarified by 10min centrifugation at 10 000rpm, 4°C. Shearing of the DNA was checked after reversing the crosslinking on agarose gel electrophoresis to be around 300-500bp. Sheared chromatin was quantified by optical density (260nm) and diluted 10-fold in IP buffer to final concentrations: 1% TRITON, 0.1% NaDeoxycholate, 0.1% SDS, 150mM NaCl, 10mM TRIS pH8, 1mM EDTA, 0.5mM EGTA, 1X protease and phosphatase inhibitors). 15µg of chromatin and 2µg of antibodies, in a final volume of 500µL, were incubated at 4°C for 16h on a wheel. 25µL of saturated magnetic beads coupled to protein G (Dynabeads) were used to recover the immuno-complexes. After 2h of incubation the bound complexes were washed extensively 5min at room temperature on a wheel in the following wash buffers: WBI (1% TRITON, 0.1% NaDOC, 150mM NaCl, 10mM TRIS pH8), WBII (1% TRITON, 1% NaDOC, 150mM KCl, 10mM TRIS pH8), WBIII (0.5% TRITON, 0.1% NaDOC, 100mM NaCl, 10mM TRIS pH8), WBIV (0.5% Igepal CA630 (Sigma), 0.5% NaDOC, 250mM LiCl, 10mM TRIS pH8, 1mM EDTA), WBV (0.1% Igepal, 150mM NaCl, 20mM TRIS pH8, 1mM EDTA), WBVI (0.001% Igepal, 10mM TRIS pH8). 20µg of sheared chromatin used as input and ChIP beads were then boiled 10min in 100µL H_2_O containing 10% (V/W) chelex resin (BioRad), followed by Proteinase K (0.5mg/mL)-digestion for 30min at 55°C, and then finally incubated 10min at 100°C. 1µL of ChIP eluate was used for qPCR assays in 10μL reactions with Brillant III Ultra Fast SYBR-Green Mix (Agilent) using a Stratagene MX3005p system. The analysis of qPCR was performed using the MxPro software. Primers are listed in **Table S1.**

### Immunoblotting

Approximately 500µm of muscle cryosections from frozen muscle were homogenized with a Dounce homogenizer protein extraction buffer containing 50mM Tris-HCl, pH 7.4, 100mM NaCl, 0,5% NP40, with Halt Protease and Phosphatase inhibitor cocktail (78440, ThermoFisher Scientific). Samples were then centrifuged for 10min at 1500g at 4°C and supernatants were collected. Protein concentration was determined by Bradford assay (ThermoFisher Scientific). Samples were denatured at room temperature for 30 min (Ca_V_β1 proteins), or at 95°C for 5min (all other proteins), with Laemmli buffer with 5% βmercaptoethanol. Proteins were separated by electrophoresis (Nu-PAGE 4–12% Bis-Tris gel; Life Technologies) and then transferred to nitrocellulose membranes (GE Healthcare). Membranes were blocked with 5% BSA (CaVβ1 proteins) or 5% milk (all other proteins) and incubated overnight at 4°C with primary antibodies (listed in **Table S3**) diluted in the blocking buffer. Membranes were then labelled with fluorescent secondary antibodies (listed in **Table S4**) diluted in blocking buffer for 1h in the dark. Images were acquired with camera LAS4000 (GE Healthcare). Western blot image analysis was performed with the public domain software ImageJ (analyze gel tool).

### Immunofluorescence and image acquisition – muscle cryosections

Immunofluorescence procedures were performed on 10µm muscles cryosections fixed on glass slides and stored at −80°C. Slides were rehydrated in PBS, fixed with PFA 4% for 10 min, permeabilized with 0.5% Triton X-100 (Sigma-Aldrich) and blocked in PBS with 4% bovine serum albumin, 0.1% Triton X-100 for 1h. Sections were incubated in PBS with 2% BSA, 0.1% Triton X-100 with primary antibodies (listed in **Table S3**) overnight at 4°C. Aspecific sites were blocked with BSA 5% 1h and slides were incubated for 1h with secondary antibodies (listed in **Table S4**) and incubated 5min with 4’,6’-diamidino-2-phenylindole (DAPI) for nuclear staining and mounted in Fluoromount (Southern Biotech).

### Immunofluorescence and image acquisition – muscle fibers

TA muscles were collected, rinsed once in PBS and fixed in a PBS 4% PFA solution at room temperature for 1 hour. Groups of approximately 10 muscle fibers were gently dissected and incubated overnight in PBS with 0.1M glycine at 4°C with mild shaking. The next day, after a day of washing in PBS 0.5% Triton X-100, fibers were incubated in permeabilized and blocked for 6 hours in PBS 3% BSA 5% goat serum 1% Triton X-100 at room temperature. Fibers were incubated with primary antibodies (**Table S3**) at 4°C for 48hin blocking solution. Fibers were then rinsed for several hours at room temperature, in PBS 0.5% Triton X-100 and incubated overnight at 4°C with secondary antibodies (**Table S4**) and α-Bungarotoxin-Alexa Fluor conjugated-594 (**Table S3**) in blocking solution. Finally, after a day of washing in PBS 0.5% Triton X-100, fibers were mounted on slides in non-hardening Vectashield mounting medium (H-1000, Vector Laboratories).

NMJ images were acquired with Zeiss Axio Observer microscope coupled with apotome module (63x objective) and edited with Zeiss Zen Lite 3.7 software. The same laser power and setting of parameters were used throughout to ensure comparability. The images presented were single projected images obtained by overlaying sets of collected z-stacks. NMJ morphometric analysis was performed on confocal z-stack projections of individual NMJs by using ImageJ-based workflow adapted from NMJ-morph1. At least 100 NMJs have been analyzed per condition. NMJ morphometric analysis was performed in a blinded manner by the same investigator.

### Immunofluorescence and image acquisition – primary myotubes

For immunofluorescence analyses, primary myotubes were seeded and cultured in fluorodishes (Dutscher). Cells were fixed in 4% PFA for 10 min, permeabilized with PBS 5% Triton X-100. Aspecific sites were blocked with BSA 1% and goat serum 10% for 30 min. Cells were incubated with primary antibodies (listed in **Table S3**) overnight at 4°C in PBS 0.1% saponin 1% BSA. Cells were washes three times and then incubated with secondary antibodies (listed in **Table S4**) and DAPI (1:10 000) for 1h in the dark.

Fluorescence images of myotubes were acquired on Spinning Disk confocal scanner unit (CSU-W1 YOKOGAWA) with a 40x oil-immersion objective (plan fluor 40x/1.3 oil OFN25 DIC N2 cyan), coupled with a Photometrics PRIM 95B Camera. Software used was MetaMorph 7.10.2. Image analyses and quantification of AChR clusters size and distribution were performed with the public domain software ImageJ by using a macro developed in the lab by MyoImage facility.

### Statistical analyses

For comparison between two groups, data were tested for normality using a Shapiro–Wilk test and for homoscedasticity using a Bartlett test followed by parametric (two-tailed paired, unpaired Student’s t-tests) or non-parametric test (Mann-Whitney) to calculate p-values. For comparison among more than two groups, according to normality ordinary one-way ANOVA or Kruskal-Wallis tests were performed. According to homoscedasticity, Brown-Forsythe ANOVA or ANOVA were performed. When it was necessary, two-way ANOVA tests were performed to compare between groups All ANOVA and Kruskal-Wallis were followed by appropriated post-hoc tests. All statistical analyses were performed with GraphPad Prism 8 or 9 software statistical significance was set at p < 0.05.

## RESULTS

### Identification of a novel Ca_V_β1 isoform in adult denervated and in embryonic skeletal muscles

The first description of the *Cacnb1E* embryonic isoform emerged from RT-PCR and genome-wide transcriptomic analyses (1). This study revealed modulation of *Cacnb1* expression at the exon level. However, the existence of several predicted though unidentified isoforms sharing the same exons, adds complexity to the interpretation of the results. To address this issue and go deeper in this characterization, we delineated the landscape of full-length Cacnb1 transcripts in adult innervated and denervated mouse skeletal muscle using targeted Oxford Nanopore sequencing (16, 17) (Tibialis Anterior: TA; Gastrocnemius: GAS), two conditions in which modulation of the expression of *Cacnb1* isoforms has already been observed (1) (**Fig 1A; Fig S1**). The visualization of the high-quality aligned Nanopore reads revealed the vast diversity of *Cacnb1* transcript variants, but with one major isoform per transcription unit (TU): more than 94% of the transcripts of the TU#1, ranging from exon1 to exon14 corresponds to *Cacnb1E* isoform and more than 83% of the transcripts of the TU#2, ranging from exon2B to exon13end correspond to *Cacnb1D* isoform. The examination of the TU#3, ranging from exon1 to exon13end unveiled that more than 93% of the transcripts of this TU correspond to *Cacnb1A* (**Fig 1A; Fig S1**). It should be noted that these percentages represent the predominance of each isoform within their respective transcriptional units, not their overall abundance in the adult muscle. We validated these data by performing RT-PCR analyses of full-length transcripts in adult innervated and denervated as well as embryonic and perinatal limb mouse muscles (1). We confirmed that these three transcripts were expressed in adult mouse muscle and that the expression of both *Cacnb1A* and *Cacnb1E* significantly increased in denervated muscle (*Cacnb1A*: P=0.0041; *Cacnb1E:* P=0.0001), while *Cacnb1D* decreased (**Fig 1B**). By amplifying full-length transcripts of *Cacnb1A*, *E* and *D* from embryonic (E), newborn (P0) and adult (P120) mouse muscle extracts, we observed the appearance of *Cacnb1D* at P0 to be fully expressed in adult as expected (**Fig 1C**). Additionally, both *Cacnb1A* and *E* transcripts were expressed during embryogenesis and adulthood (**Fig 1C**). We analyzed protein expression of Ca_V_β1 isoforms by western blot using a custom antiserum antibody specific to Ca_V_β1D and A variants (epitope coded by RNA sequence in exon 13end). However, for Ca_V_β1E no specific antibody for this isoform could be found, neither customized nor commercially available. In adult muscle, the levels of a 58kDa band, corresponding to Ca_V_β1A, increased in denervated muscle, while the 53kDa band intensity, corresponding to the Ca_V_β1D, did not change in this condition (**Fig 1D**). Wondering about the expression kinetic of Ca_V_β1D and Ca_V_β1A variants at various age stages, we measured their levels in embryonic muscles at E12.5 and E16, and at post-natal days P0, −7, −14 and − 30 by using the same custom antibody. We confirmed that Ca_V_βA appeared in E16 until P14 and became weakly expressed in P30 innervated muscle whereas Ca_V_βD started at P0 to be stably expressed in adult muscle **(Fig1E).**

**Figure 1.**
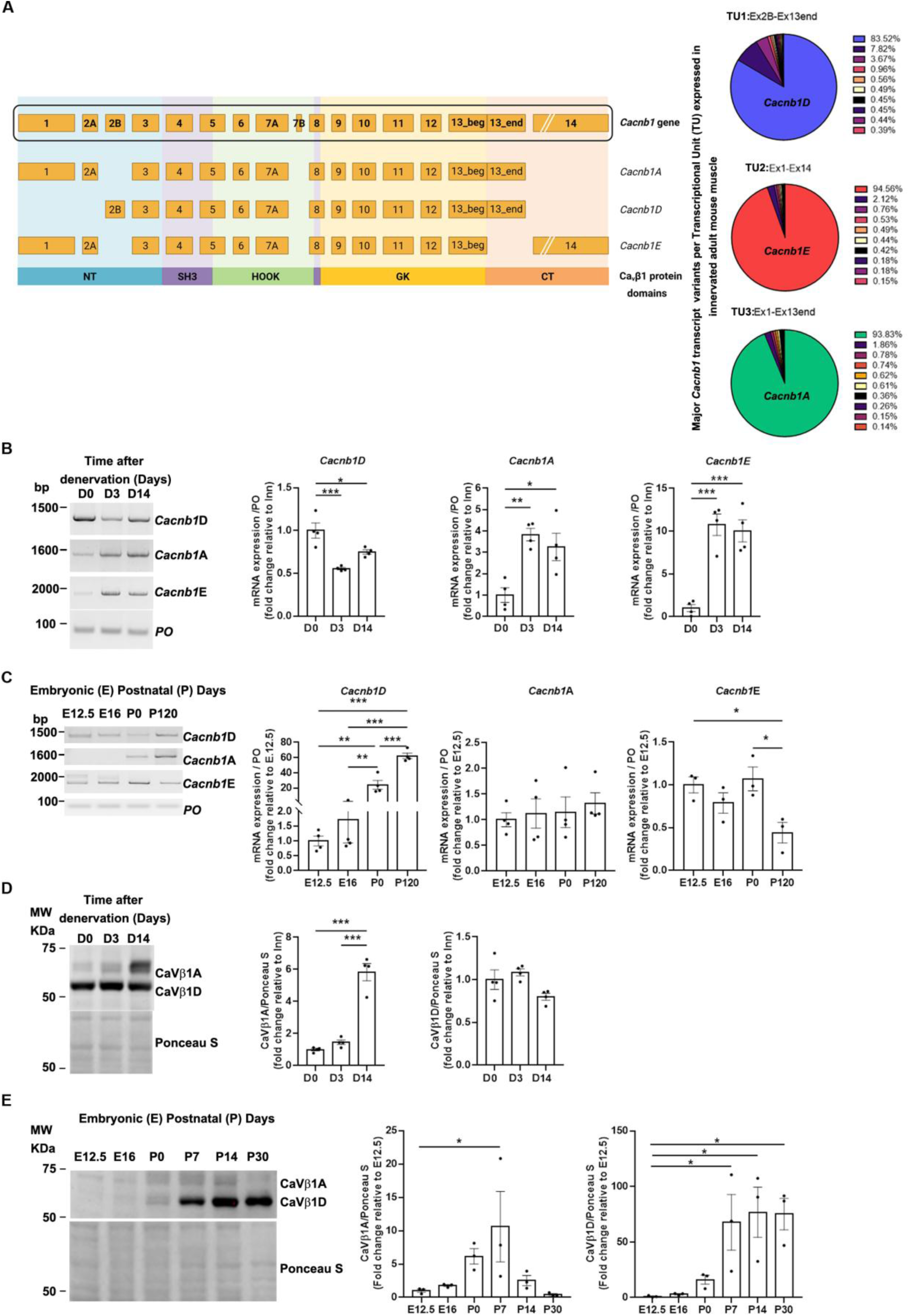
Identification of a new Ca_V_β1 isoform in embryonic muscles. **A.** *Cacnb1* gene and skeletal muscle transcript variants: *Cacnb1A, Cacnb1D* and *Cacnb1E* as well as the corresponding protein domains are represented. Full-length sequencing of *Cacnb1* transcript through Nanopore technology led to the identification of one major isoform per TU. **B.** RT-PCR and quantification of the three full-length *Cacnb1* variants in innervated (D0) TA muscles or after 3 (D3) and 14 days of denervation. PO was the loading control. **C.** Western Blot and quantification of Ca_V_β1A and Ca_V_β1D in embryonic (E) and postnatal (P) days. Ponceau S was the loading control. **D.** Western Blot and quantification of Ca_V_β1A and Ca_V_β1D after 3 (D3) and 14 days of denervation. **E.** Western Blot and quantification of Ca_V_β1A and Ca_V_β1D in embryonic (E) muscles and adult (P) TA muscles. Ponceau S was the loading control. All data are mean ± SEM (\**P* < 0.05, \*\**P* < 0.01, \*\*\**P* < 0.001). *P* values were calculated by ordinary one-way ANOVA followed by (B-D) Tukey’s and (E) Dunnett’s multiple comparison test.

Overall, our results reveal that *Cacnb1A*, and not only *Cacnb1E,* transcripts are expressed in mouse skeletal muscle during embryogenesis until first post-natal life and increases after nerve damage, with Ca_V_β1A protein following this pattern, whereas Ca_V_β1E, due to the demission of previously commercialized antibodies, cannot be detected at protein level. In addition, we validate the expression of *Cacnb1D* transcript and the corresponding Ca_V_β1D protein exclusively after birth. Taken together, these data underline the complexity of Ca_V_β1 isoforms and intrigue about the specific role of each of these Ca_V_β1 proteins in skeletal muscle physiology.

### Activation of distinct promoters for *Cacnb1* isoforms expression in embryonic and adult muscles

To further investigate the transition of embryonic versus adult Ca_V_β1 isoforms, we performed triplex RT-PCR analyses for deciphering the kinetic of exon2B inclusion that characterizes the adult *Cacnb1D* isoform, versus the kinetic of exon2A exclusion for identifying the embryonic *Cacnb1A/E* variants. As expected, we observed that exon2A was present from E12.5, while exon2B expression increased at birth and was highly expressed in adult muscle (**Fig 2A**). We then wondered if *Cacnb1D* and *Cacnb1A/E* expressions might be orchestrated by the activation of distinct promoters in embryonic and in adult muscles. Nanopore sequencing revealed that the mRNAs starting at exon1 were never associated with exon2B (**Fig S1**). This suggests that the expression of exon2B may be exclusively triggered by a promoter located within this exon. To validate this hypothesis, we performed an *in silico* analysis (ENCODE project, UCSC Genome browser) of ChIPseq signals of methylation and acetylation histone marks and RNA-Polymerase II occupancy on mouse *Cacnb1* gene in embryonic muscles compared to adult hind limb muscles. We found that methylation marks H3K4me3, H3K9ac and H3K27ac, characteristic of active promoters (18), were mostly located at exon1 in E12.5 mouse muscle, while H3K27ac was mainly located at exon2B in three different adult mouse hind limb muscles (**Fig 2B**). In addition, RNA-Polymerase II was recruited at *Cacnb1* exon1 in E12.5 mouse muscle and at exon2B in adult mouse muscle (**Fig 2B**), suggesting a transcription initiation site (19). Together, these data strongly suggest that distinct and specific promoters are used for the expression of embryonic (Prom1) versus adult (Prom2) *Cacnb1* isoforms. We then asked whether the innervation status in adult muscle could affect the activity of *Cacnb1* promoters, therefore modulating the expression of *Cacnb1* isoforms. We found that the percentage of exon2B inclusion was higher in innervated and decreased in denervated muscles whereas exon1 inclusion was very low in innervated and increased significantly in denervated muscle RNA extracts (**Fig 2C**). To evaluate the activity of the putative specific promoters, we performed a ChIP-PCR analysis on *Cacnb1* gene. We observed a reduction in epigenetic marks of active promoters (H3K4me3 and H3K9ac) at Prom2 site, while these marks were enriched around Prom1 following denervation. Accordingly, the quantification of RNA-Polymerase II recruitment displayed increased enzyme’s occupancy around Prom1 and decreased enzyme’s occupancy at Prom2 site in denervated compared to innervated muscles (**Fig 2D, Fig S2**).

**Figure 2.**
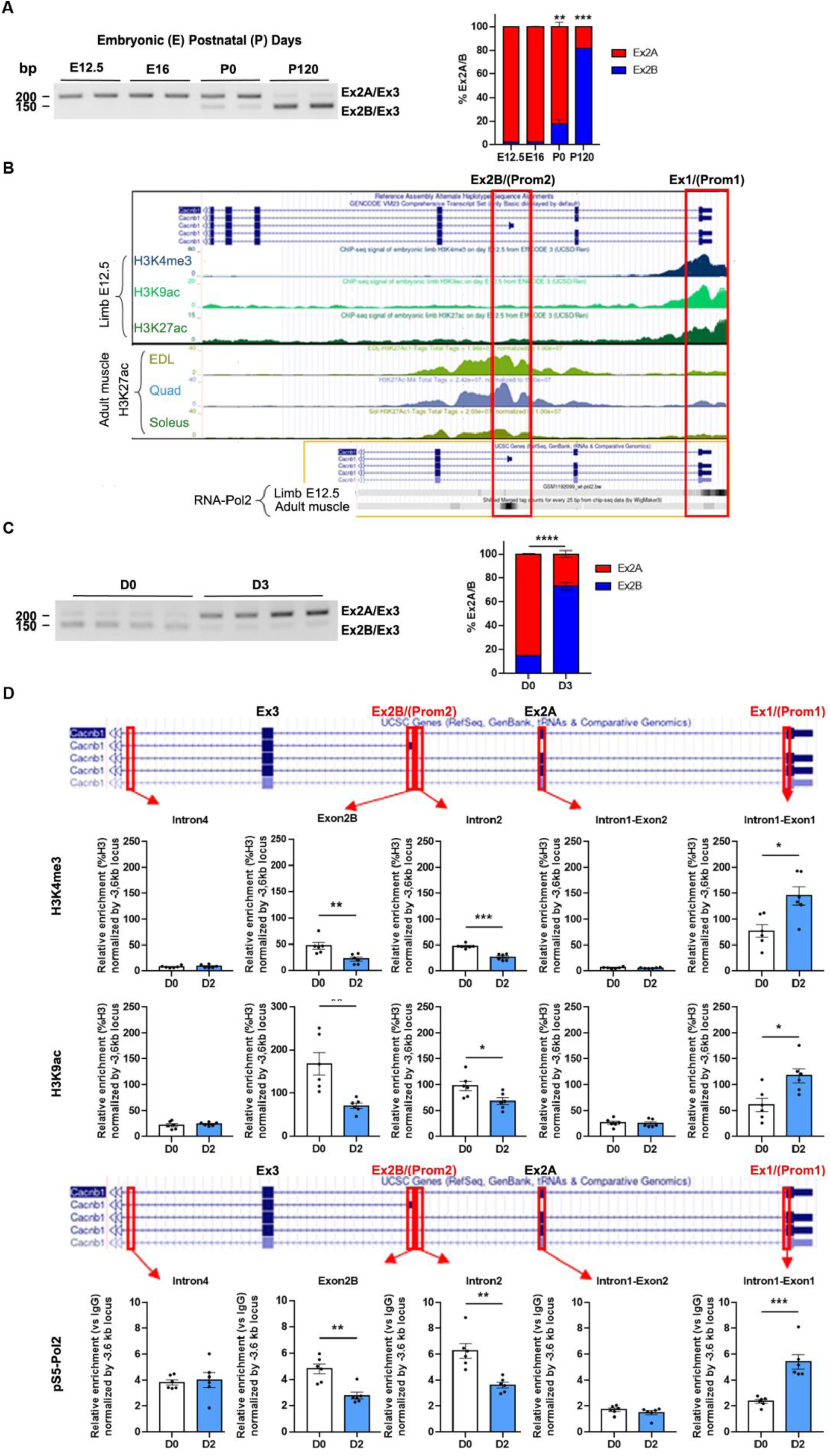
Two specific and distinct promoters drive the expression of *Cacnb1* isoforms in skeletal muscle. **A.** Triplex RT-PCR and quantification of the percentage of exon2A versus exon2B inclusion in embryonic (E) and adult (P) TA muscles. **B.** Visualization of epigenetic marks and RNA Polymerase II at *Cacnb1* promoters using UCSC Genome Browser. Image displaying the location of exon 2B (Ex2B) - containing the promoter site 2 (Prom2) -, exon 1 (Ex1) containing the promoter site 1 (Prom1), activating epigenetic marks (H3K3me3, H3K9ac and H3K27ac) and RNA Polymerase II occupancy at Prom1 and Prom2 in either embryonic (E12.5) muscles (ENCODE project) or adult muscles (Extensor digitorum longus (EDL), Quadriceps and Soleus). **C.** Triplex RT-PCR and quantification of the percentage of exon2A versus exon2B inclusion in innervated and 3-days denervated adult TA muscles (D3). **D.** Chromatin immunoprecipitation (ChIP) followed by PCR analysis showing the presence of activating epigenetic marks (H3K4me3 and H3K9ac) and Serine5-phosphorylated RNA Polymerase II (pS5-Pol2) occupancy at Prom1 and Prom2 in innervated and 2-days denervated adult TA/Gas muscles. All data are mean ± SEM (\**P* < 0.05, \*\**P* < 0.01, \*\*\**P* < 0.001). *P* values were calculated by ordinary one-way ANOVA followed by Tukey’s multiple comparison test (**A**) or unpaired t-test (**C**, **D**).

Taken together, our results decipher the *Cacnb1* isoform transition from embryonic to adult isoforms and show that its regulation is driven by the activation of two distinct and specific promoters. Furthermore, we demonstrate that adult skeletal muscle restores the embryonic epigenetic program in *Cacnb1* gene after nerve damage, suggesting a role for *Cacnb1* in the mechanisms aiming at recovering or maintaining a proper innervation status and potentially in other events associated with neuromuscular physiology.

### Role of embryonic Ca_V_β1 isoforms in AChR clustering

The expression of different *Cacnb1* isoforms at embryonic and adult stages has raised the question of the specific role of different variants in the formation, maturation and maintenance of the NMJ.

We investigated the role of embryonic *Cacnb1* isoforms in agrin-mediated AChR clustering in highly differentiated primary myotubes (13, 20, 21) by downregulating these isoforms using an adeno-associated virus 1 (AAV1) carrying a short hairpin targeting a sequence expressed in exon2A of *Cacnb1* pre-RNA, thus silencing specifically Ca_V_β1A and E transcripts and proteins (shEx2A) (1) (**Fig 3A**). After validating their downregulation (**Fig 3A**), we observed that their absence had no effect on AChR aggregation capacity, showed by a similar number of AChR clusters per myofiber surface compared to the control condition (**Fig 3B, C**). However, it induced a significant raise of AChR cluster size, with a mean fold increase of 2.93 (**Fig 3B, C**). Additionally, we observed an augmented distance between AChR aggregates and the nearest nuclei, suggesting that embryonic Ca_V_β1 isoforms may play a role in positioning the future sub-synaptic nuclei (**Fig 3B, C**). To identify the mechanisms responsible for these *in vitro* AChR clustering abnormalities, we measured protein expression of AChRα as well as key players in aggregate formation including MuSK and Dok7 (9, 22, 23). Although we did not observe a significant modification of AChRα and Dok7 protein levels, we found higher MuSK expression after silencing Ca_V_β1A and E compared to control (**Fig 3D**).

**Figure 3.**
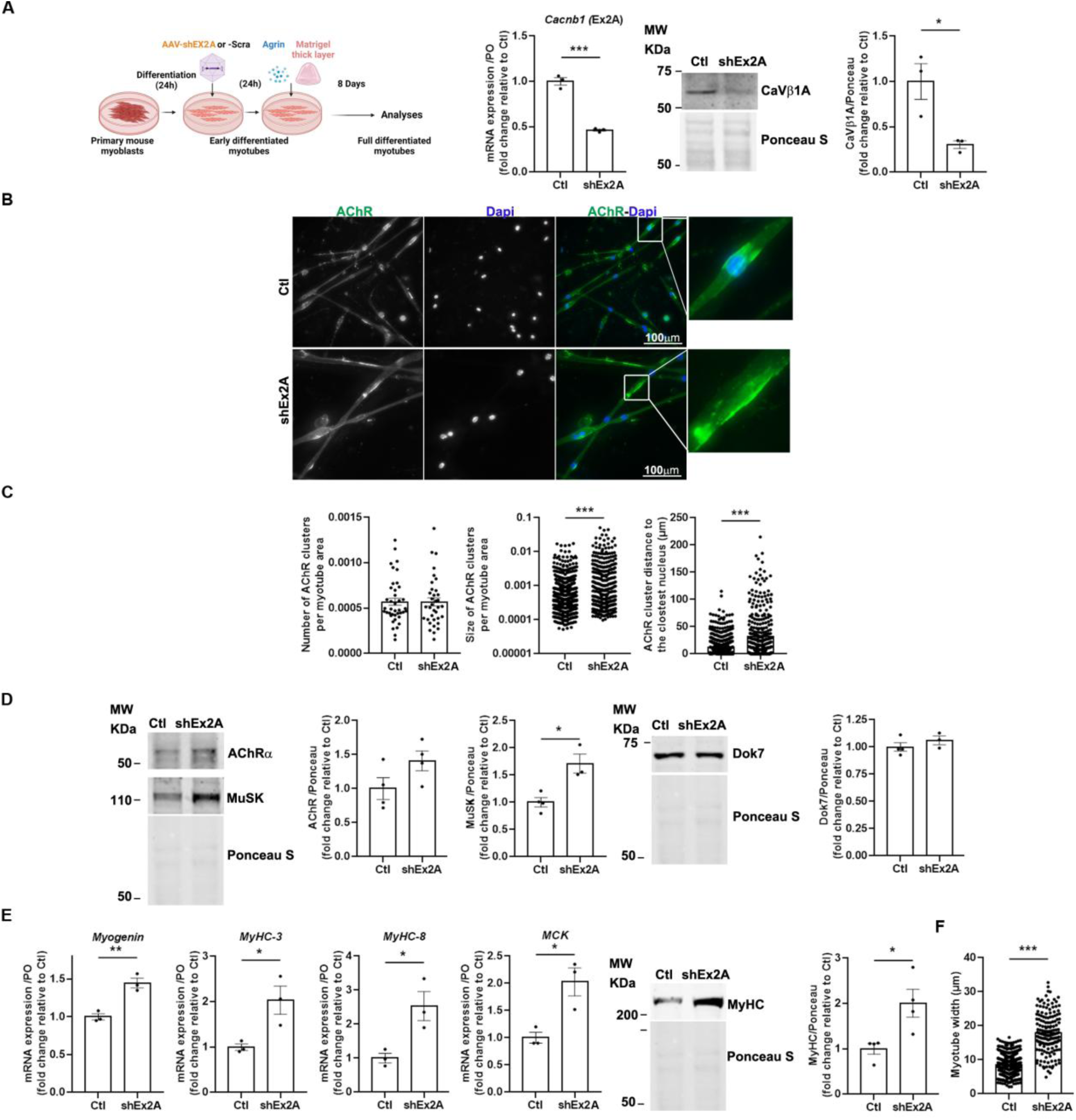
Downregulation of embryonic Ca_V_β1 isoforms induces bigger AChR aggregates and myotube precocious maturation in an *in vitro* model of highly differentiated myotubes. **A.** Schematic representation of the experimental protocol. RT-qPCR of *Cacnb1* exon 2A (Ex2A), Western Blot and quantification of Ca_V_β1A in control (Ctl) and shEx2 treated myotubes. **B.** Immunofluorescence (IF) images of control or shEx2A treated myotubes stained with AChRa1 (green) and DAPI (blue). **C.** Quantification of AChR clusters number and size per myotube area and distance between AChR clusters and the nearest nuclei. **D.** Western Blot and quantification of AChR and MuSK in control and shEx2A treated myotubes. **E.** RT-qPCR of *Myogenin*, *MyHC-3*, *MyHC-8* and *MCK* in control and shEx2A treated myotubes. Western Blot and quantification of MyHC in control and shEx2A treated myotubes. All data are mean ± SEM (\**P* < 0.05, \*\**P* < 0.01, \*\*\**P* < 0.001). *P* values were calculated by unpaired t-test (**C**, **D**).

Then, we investigated the potential role of embryonic *Cacnb1* isoforms on the myogenic differentiation process. Despite enhanced *Myogenin* expression, we measured significantly higher *MyHC*-3, *MyHC*-8 and *MCK* mRNA expressions (p=0.0318, p=0.027 and p=0.0203 respectively), as well as MyHC protein expression levels in shEx2A myotubes compared to controls (**Fig 3E**). These data suggest that myogenic maturation was aberrantly accelerated in the absence of embryonic *Cacnb1* isoforms, corroborated by the finding that shEx2A treated myotubes were wider compared to controls (**Fig 3F**). Altogether, these findings indicate that embryonic/perinatal Ca_V_β1 isoforms are needed not only to modulate the first phases of myogenesis as previously shown (3), but also to ensure the coordinated terminal muscle cell differentiation and AChR aggregation in response to agrin stimulation.

### Role of embryonic Ca_V_β1 isoforms in post-natal NMJ maturation/maintenance

The presence of embryonic/perinatal Ca_V_β1 isoforms expressed until at least P14 (**Fig 1E**), suggests their possible involvement in NMJ postnatal maturation/maintenance. To decipher the role of these proteins in that process, we downregulated their expression using AAV-shEx2A injected in the hind limb muscles of 7-days-old pups (**Fig 4A**), and analysed NMJ morphology 21 days later (P30 mice), when endplates are usually mature (24). We quantified NMJ morphological parameters and found that the AChR area was significantly reduced upon embryonic *Cacnb1* variants depletion compared to control, while the size of AChR area unoccupied by nerve endings was increased (**Fig 4B, C**). Additionally, the overlap between nerve terminals and post-synaptic apparatus was decreased and a significant NMJ fragmentation was observed (**Fig 4B, C**). Although NMJ morphology appeared altered with pre and postsynaptic abnormalities, we did not observe alterations in AChRα, MuSK, Dok7 protein expression (**Fig 4D)** nor in *Myogenin* gene expression **(Fig 4E).**

**Figure 4.**
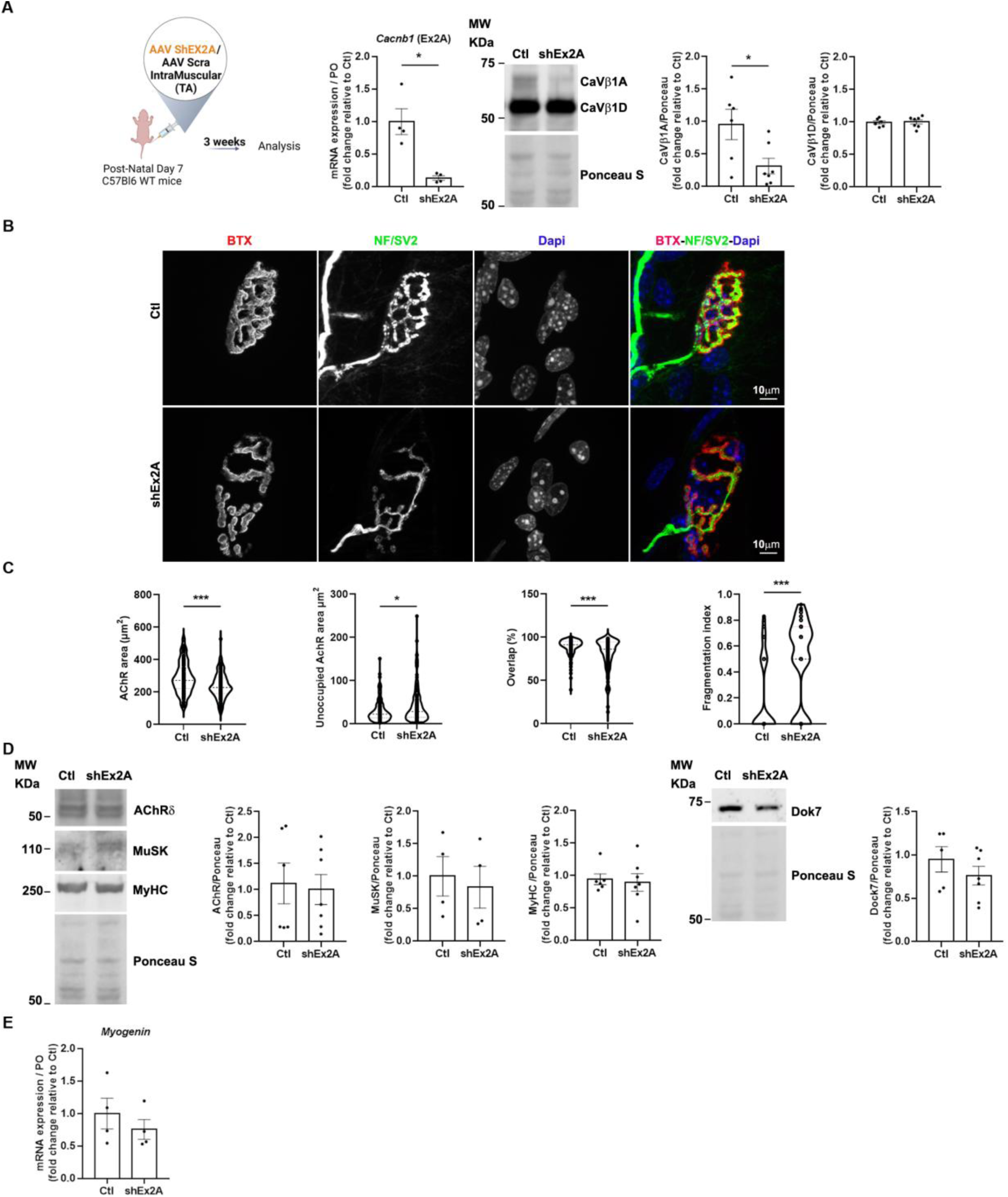
NMJ structural and molecular defects upon downregulation of embryonic Ca_V_β1 isoforms in TA muscles at postnatal stages. **A.** Schematic representation of the experimental protocol. RT-qPCR of embryonic *Cacnb1* variants using primer in exon2A in control and shEx2 treated TA muscles, Western Blot and quantification of Ca_V_β1A and Ca_V_β1D in muscles 3-weeks post-injection. **B.** Immunofluorescence (IF) images of control and shEx2A treated TA muscles, 3-weeks post-injection, stained with α-bungarotoxin (BTX) for AChR (red), neurofilaments and Synaptic Vesicle Glycoprotein 2 (NF/SV2) (green) and DAPI (blue). **C.** Quantification of NMJ morphology with area of AChR, unoccupied AChR area, overlap between nerve terminals and post-synaptic apparatus and fragmentation index. **D.** Western Blot and quantification of AChRδ, MuSK and MyHC in control and shEx2A treated TA muscles, 3-weeks post-injection. **E.** RT-qPCR of *Myogenin*, in control and shEx2A treated muscles. All data are mean ± SEM (\**P* < 0.05, \*\**P* < 0.01, \*\*\**P* < 0.001). *P* values were calculated by paired (**A –** *Cacnb1* RT-qPCR-, **E**) and unpaired t-test (**A**-Ca_V_β1A and Ca_V_β1D WB-, **C, D**).

Overall these observations, suggest that the loss of embryonic Ca_V_β1 isoforms during early post-natal stages significantly impacts endplate morphology without inducing major abnormalities in the expression of key synaptic proteins involved in its formation nor perturbation of myofiber maturation.

### Role of embryonic Ca_V_β1 isoforms in NMJ maintenance in adult skeletal muscle

To better understand if Ca_V_β1A and Ca_V_β1E isoforms may play a role in the stabilization of nerve-muscle synapses once they are completely formed and mature, we disrupted their expression in TA of adult mice via the AAV-shEx2A injection (**Fig 5A**). We then analysed morphological, molecular and functional readouts reflecting NMJ integrity 12 weeks post-injection. The quantification of pre and post synaptic morphological parameters revealed that the perimeters and areas of both AChR and endplates were significantly reduced upon embryonic *Cacnb1* depletion compared to controls. Additionally, the size of synaptic contact and the overlap between nerve terminals and post-synaptic apparatus were decreased. Furthermore a significant NMJ fragmentation was observed upon downregulation of embryonic Ca_V_β1 proteins **(Fig 5B, C).**

**Figure 5.**
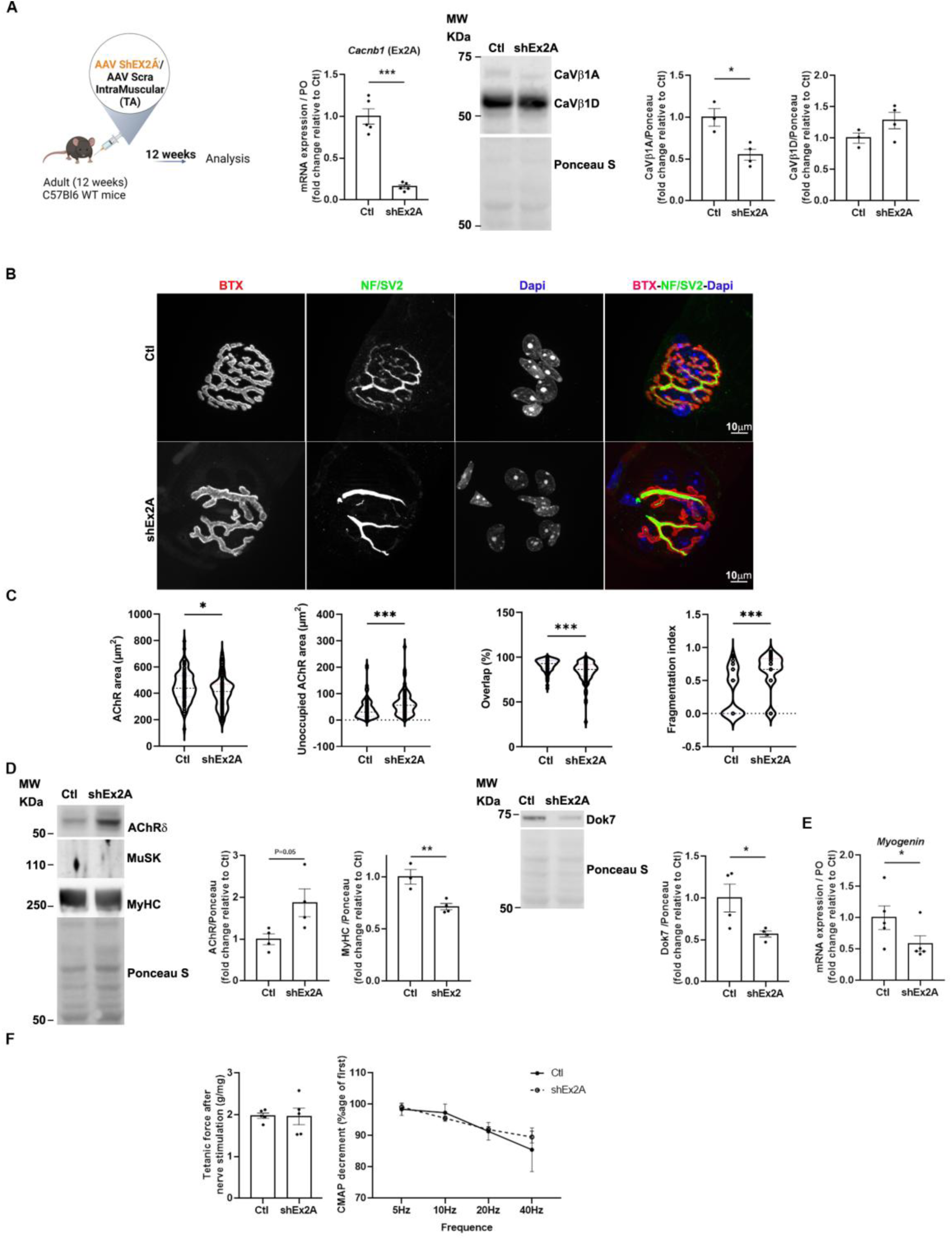
NMJ transcriptional and functional defects upon downregulation of embryonic Ca_V_β1 isoforms in adult TA muscles. **A.** Schematic representation of the experimental protocol. RT-qPCR of embryonic *Cacnb1* variants using primer in exon2A in control and shEx2A treated TA muscles, Western Blot and quantification of Ca_V_β1A and Ca_V_β1D in muscles 12-weeks post-injection. **B.** Immunofluorescence (IF) images of control and shEx2 treated TA muscles, 12-weeks post-injection, stained with α-bungarotoxin (BTX) for AChR (red), neurofilaments light chain and Synaptic Vesicle Glycoprotein 2 (NF/SV2) (green) and DAPI (blue). **C.** Quantification of NMJ morphology with area of AChR, unoccupied AChR area, overlap between nerve terminals and post-synaptic apparatus and fragmentation index. **D.** Western Blot and quantification of AChRδ, MuSK and MyHC, of Dok7, in control and shEx2A treated TA muscles 12-weeks post-injection. **E.** RT-qPCR of *Myogenin*, in control and shEx2A treated muscles 12-weeks post-injection. **F.** Indirect (nerve) or direct (muscle) tetanic force measured in control and shEx2A treated TA adult muscles, 12-weeks post-injection. CMAP decrement measured by ENMG in control and shEx2A treated TA adult muscles, 12-weeks post-injection. All data are mean ± SEM (\**P* < 0.05, \*\**P* < 0.01, \*\*\**P* < 0.001). *P* values were calculated by or paired (**A –** *Cacnb1* RT-qPCR-, **E, F**) or unpaired t-test (**A**-Ca_V_β1A and Ca_V_β1D WB-, **C, D**).

Analysis of molecular players involved in NMJ formation and maintenance highlighted a significant increase in AChRδ protein and transcript expression, whereas the expression of MuSK protein was undetectable in all conditions (**Fig 5D**). In addition, we observed significant decrease in Dok7 and MyHC protein expression **(Fig 5D),** and a decrease in *Myogenin* transcript levels (**Fig 5E**). Next, we measured muscle function in terms of force generation, by direct (muscle) and indirect (nerve) stimulation and the neuromuscular connectivity by electroneuromyogram (ENMG) measurement of compound muscle activity potential (CMAP) decrement. We found that the ablation of embryonic Ca_V_β1 proteins did not affect all these parameters, indicating that NMJ morphological alterations had no impact on muscle function, at least in that observed time window.

Altogether, these data suggest that embryonic/perinatal Ca_V_β1 isoforms are 1) likely involved in NMJ formation prior to innervation, as supported by the *in vitro* study, 2) involved in the maintenance and/or maturation of the post-synaptic apparatus at early postnatal stages, and 3) involved in NMJ maintenance in adult muscles.

## DISCUSSION

Neuromuscular system plays a vital role for all vertebrates. Its proper development, maturation and maintenance are strictly regulated processes requiring a multitude of mechanisms not yet completely elucidated. In this context, some years ago it has been described, that a component of ECC machinery, Ca_V_β1, the regulatory subunit of muscle Voltage-gated Ca^2+^ channel encoded by the *Cacnb1* gene, was essential in the formation of nerve-muscle pre-patterning during embryonic development (5). In a further study this protein has been demonstrated as having transcription factor properties regulating myogenic factor in muscle precursor cells (3). More recently, two Ca_V_β1 skeletal muscle isoforms have been identified by our work, Ca_V_β1D, defined as the adult constitutive isoform and Ca_V_β1E defined as an embryonic isoform with a role in up-regulating the expression of *Gdf5* and thus limiting muscle mass loss after denervation and during aging (1).

Here, we aimed to further characterize Ca_V_β1 isoforms by investigating the mechanisms behind the regulation of their expression, and deciphering their role in the formation, maturation and stability of neuromuscular system. The use of Nanopore sequencing as innovative and powerful exploring technology, allowed identification of *Cacnb1A*, coding for Ca_V_β1A protein, an additional embryonic isoform, expressed in adult muscle after denervation. Investigating the mechanisms regulating the differential expression of *Cacnb1* transcripts, we have discovered that adult *Cacnb1D* and embryonic/perinatal *Cacnb1A* and *Cacnb1E* are expressed through the activation of two distinct promoters located in exon1 (Prom1) and exon2B (Prom2), which are differently regulated during development. Indeed, we have demonstrated that Prom1 is active during embryogenesis, while Prom2 is active in mature adult muscle, perinatal stages marking a period of transition between the activation of these two promoters. In addition, we have shown that, in adult muscle, nerve damage leads to the restoration of the embryonic epigenetic program of *Cacnb1* promoters. These evidences shed light on the molecular events behind Ca_V_β1 modulation and open new hypotheses on other upstream factors potentially implicated in the fine tuning of their epigenetic program.

To gain insights into transcription factors binding at these promoter regions and potentially regulating them, we used the UCSC Genome Browser, an online software allowing comprehensive visualization and analysis of genomic data, including features like transcription factors binding sites. Among the wide range of transcription factors potentially binding to Prom1 and Prom2 regions, we identified Myogenin a muscle-specific factor known to promote transcription of genes significant for myogenesis and AChR clustering (25, 26). In addition, we found as potential binding factors of Prom2 region BRD4 and MEF2A, known to act as a regulator of catabolic and myogenesis-related genes (27, 28) and to control muscle gene transcription (29), respectively. On the other hand, we identified, as potentially binding Prom1, KDM1A (or LSD1), also described to regulate key myogenic transcription factor and modulating muscle regeneration and recovery after injury (30). More investigations are needed to define molecular interactors affecting the expression of *Cacnb1* variants, however the data presented in this study pinpointed some possible players having role in this event.

Overall, the existence of various data Ca_V_β1 variants raises further questions about the significance of multiple isoforms during development and for skeletal muscle homeostasis, especially in a VGCC-independent context. We thus investigated the role of Ca_V_β1 on the different stages on NMJ formation, maturation and stability. As mentioned, a previous study demonstrated that genetically ablated *Cacnb1* mice exhibited muscle patterning defects and aberrant innervation at E14.5 and E18.5 in mice (5). However, these alterations were observed after motor growth cones had already reached the muscle. To better understand the potential role of CaVβ1 isoforms in AChR pre-patterning *prior* to innervation—a gap that remains poorly explored in the literature—we investigated this process using an *in vitro* myotube model.

We found increased size of aneural AChR clusters associated with precocious myotube maturation, which aligns with Chen and colleagues’ post-innervation results that showed increased endplate areas and more perforated clusters, indicative of a precocious maturation. Our findings suggest that the defects observed in their study originate from abnormalities occurring before innervation begins. However, it cannot be excluded that these alterations could have been modified or amplified by nerve-derived factors.

The observed rise in AChR clusters size may be the consequence of different outcomes: i) aberrant myotube maturation that could affect clusters enlargement due to increased myotube width; ii) the increase in MuSK level, possibly resulting from a transcriptional effect of myogenin (26), may be connected with the expanded AChR clusters size (31). Further analyses would be needed to elucidate all the mechanisms governed by Ca_V_β1A and Ca_V_β1E and implicated in the orchestration of AChR pre-patterning. However, this study indicates a crucial involvement of these proteins in the first steps of NMJ formation, independently of innervation and ECC modulation.

Going further in exploring the function of Ca_V_β1A/E isoforms on endplate maturation/stability during early post-natal life, we found that ablation of these proteins in muscle of 1-month-old pups affected NMJ stricture. The observed defects could be due to alterations in maturation, stability or a combination of both and distinguishing between these two processes is a challenging purpose. Regarding the possible cause of the observed defects, the increased muscle maturation occurring under ablation of the embryonic/perinatal Ca_V_β1 isoforms could lead to NMJ fragmentation as a mechanism to extend the AChR area following precocious muscle growth. Indeed, the link between NMJ fragmentation and degeneration might not be systematic and some studies suggest that this event could also be a sign of regeneration, as a process of synaptic plasticity aimed at enlarging synaptic area (32–34).

In adult muscle, despite the apparently weak expression of Ca_V_β1A/E, their ablation lead to significant alteration of NMJ morphology. Noteworthy, similar features are observed in pathophysiological conditions as during ageing (1). Indeed, age-related morphological defects of endplates, including postsynaptic fragmentation, reduced AChR density associated with synaptic decoupling, are reported as preceding changes of NMJ molecular markers expression (35, 36). This appears to be what we observed upon Ca_V_β1A/E ablation with no global modification in NMJ-associated actors despite morphological alterations. Deeper analyses will be needed to establish if Ca_V_β1A/E downregulation in adult healthy muscle could induce a premature aging phenotype in mice, as suggested by our findings.

Taken together, our study sheds light on the effects of both Ca_V_β1A and Ca_V_β1E isoforms on distinct NMJ specific stages. It would be of high relevance in future investigations to clarify whether one or both Ca_V_β1A and Ca_V_β1E isoforms are responsible for the observed effects.

In conclusion, in this work we identified Ca_V_β1A as a novel Ca_V_β1 isoform and highlighted molecular and functional aspects of the different Ca_V_β1s as molecular players involved in the neuromuscular system formation, maturation and maintenance.

## Supporting information

list of Primers and antibodies

---

**Figure S1.**
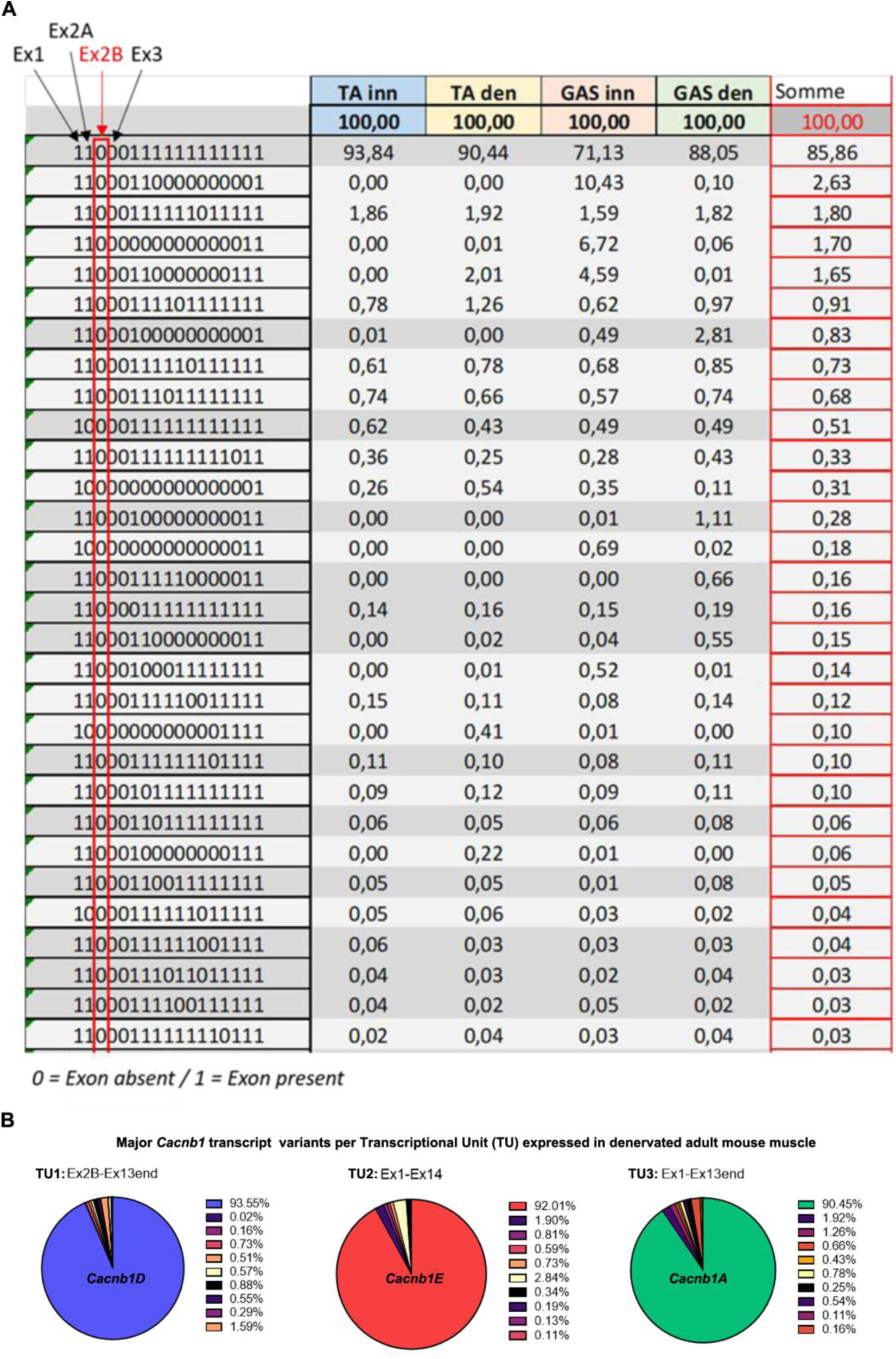
*Cacnb1* exon2B is systematically excluded when exon1 is expressed. **A.** Long-read Nanopore sequencing data depicting exon presence and absence for each *Cacnb1* variant (only the first 30 are showed) in innervated (inn) and denervated (den) mouse *Tibialis Anterior* (TA) and *Gastrocnemius* (GAS) muscles. Each row represents an individual transcript, with “1” indicating the presence and “0” indicating the absence of specific exons. This binary representation allows the observation that none of the transcripts starting at exon1 include exon2B, suggesting that alternative splicing is not the mechanism regulating exon2B inclusion but rather indicates the existence of an alternative promoter at exon 2B. **B.** Percentage of *Cacnb1* major isoform per TU after Nanopore sequencing in denervated muscle.

**Figure S2.**
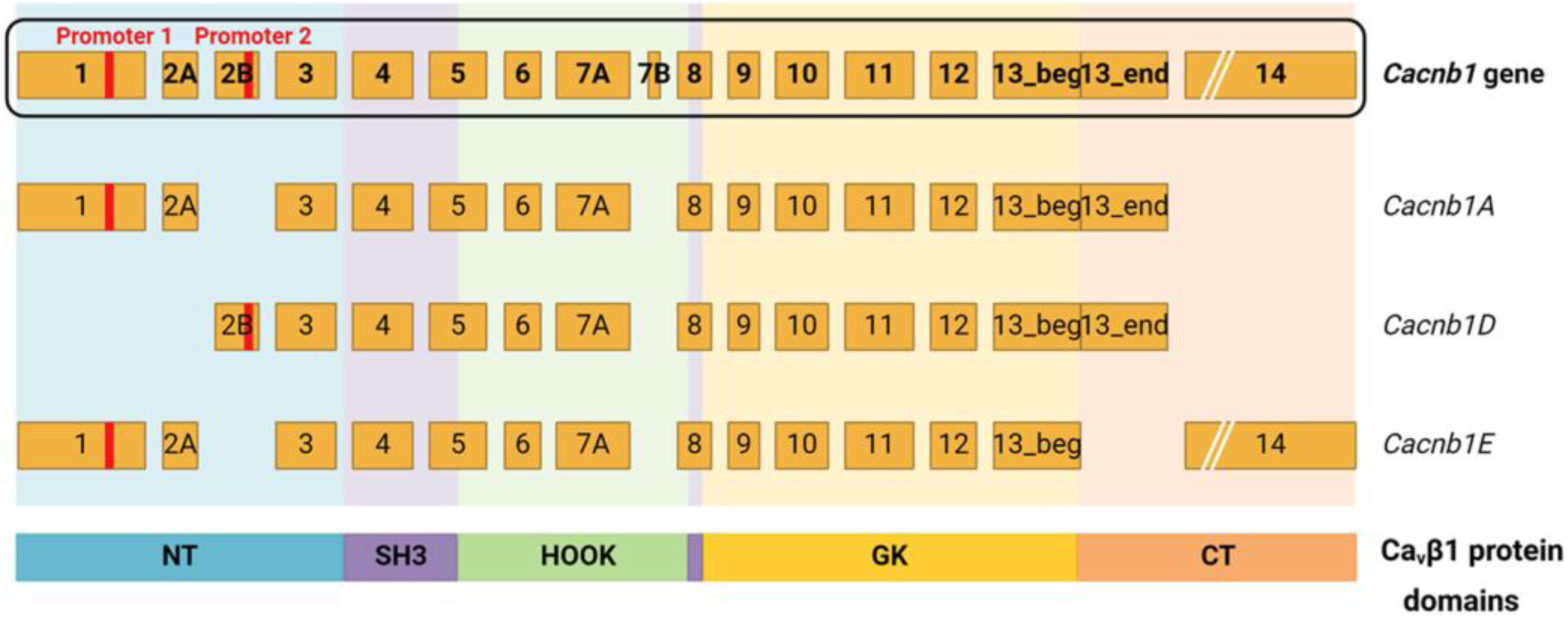
Two specific and distinct promoters drive the expression of *Cacnb1* isoforms in skeletal muscle. *Cacnb1* gene and skeletal muscle transcript variants: *Cacnb1A, Cacnb1D* and *Cacnb1E* and the corresponding promoters’ sites, as well as the corresponding protein domains are represented.

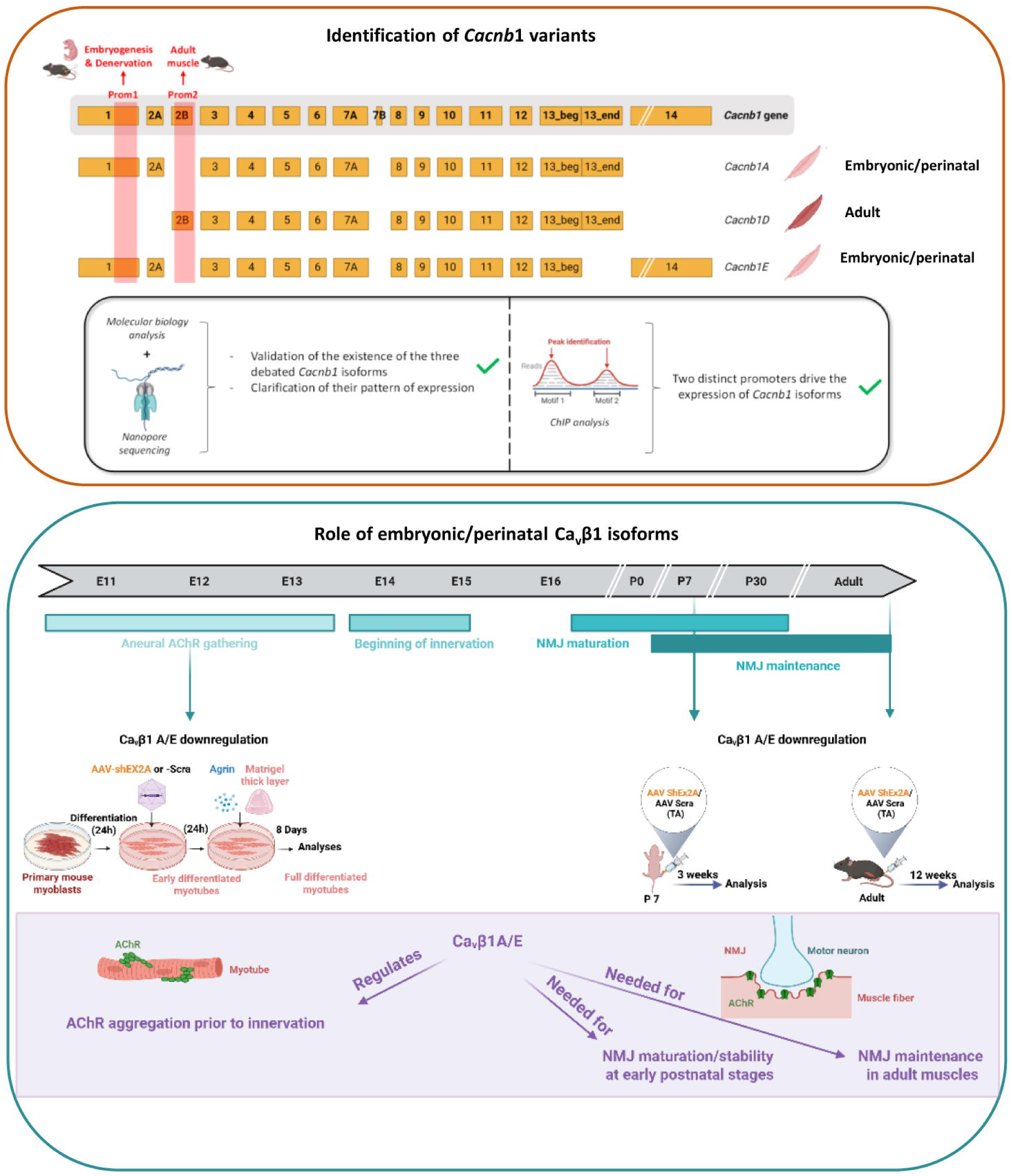

## REFERENCES

1. Traoré, M., Gentil, C., Benedetto, C., Hogrel, J.-Y., De la Grange, P., Cadot, B., Benkhelifa-Ziyyat, S., Julien, L., Lemaitre, M., Ferry, A., et al. (2019) An embryonic CaVβ1 isoform promotes muscle mass maintenance via GDF5 signaling in adult mouse. Sci Transl Med, 11, eaaw1131.

2. Buraei, Z. and Yang, J. (2013) Structure and function of the β subunit of voltage-gated Ca^2+^ channels. Biochim Biophys Acta, 1828, 1530–1540.

3. Taylor, J., Pereyra, A., Zhang, T., Messi, M.L., Wang, Z.-M., Hereñú, C., Kuan, P.-F. and Delbono, O. (2014) The Cavβ1a subunit regulates gene expression and suppresses myogenin in muscle progenitor cells. J Cell Biol, 205, 829–846.

4. Vergnol, A., Traoré, M., Pietri-Rouxel, F. and Falcone, S. (2022) New Insights in CaVβ Subunits: Role in the Regulation of Gene Expression and Cellular Homeostasis. Frontiers in Cell and Developmental Biology, 10.

5. Chen, F., Liu, Y., Sugiura, Y., Allen, P.D., Gregg, R.G. and Lin, W. (2011) Neuromuscular synaptic patterning requires the function of skeletal muscle dihydropyridine receptors. Nat Neurosci, 14, 570–577.

6. Arber, S., Burden, S.J. and Harris, A.J. (2002) Patterning of skeletal muscle. Curr Opin Neurobiol, 12, 100–103.

7. Yang, X., Arber, S., William, C., Li, L., Tanabe, Y., Jessell, T.M., Birchmeier, C. and Burden, S.J. (2001) Patterning of Muscle Acetylcholine Receptor Gene Expression in the Absence of Motor Innervation. Neuron, 30, 399–410.

8. Li, L., Xiong, W.-C. and Mei, L. (2018) Neuromuscular Junction Formation, Aging, and Disorders. Annual Review of Physiology, 80, 159–188.

9. Eguchi, T., Tezuka, T., Miyoshi, S. and Yamanashi, Y. (2016) Postnatal knockdown of dok-7 gene expression in mice causes structural defects in neuromuscular synapses and myasthenic pathology. Genes Cells, 21, 670–676.

10. Rieger, F., Powell, J.A. and Pinçon-Raymond, M. (1984) Extensive nerve overgrowth and paucity of the tailed asymmetric form (16 S) of acetylcholinesterase in the developing skeletal neuromuscular system of the dysgenic (mdgmdg) mouse. Developmental Biology, 101, 181– 191.

11. Kaplan, M.M., Sultana, N., Benedetti, A., Obermair, G.J., Linde, N.F., Papadopoulos, S., Dayal, A., Grabner, M. and Flucher, B.E. (2018) Calcium Influx and Release Cooperatively Regulate AChR Patterning and Motor Axon Outgrowth during Neuromuscular Junction Formation. Cell Rep, 23, 3891–3904.

12. Massiré, T., Chiara, N., Amélie, V., Christel, G., Marius, H., Lucile, S., Maxime, G., Anne, F., Mégane, L., Zoheir, G., et al. (2024) GDF5 as a rejuvenating treatment for age-related neuromuscular failure. Brain, 10.1093/brain/awae107.

13. Falcone, S., Roman, W., Hnia, K., Gache, V., Didier, N., Lainé, J., Auradé, F., Marty, I., Nishino, I., Charlet-Berguerand, N., et al. (2014) N-WASP is required for Amphiphysin-2/BIN1-dependent nuclear positioning and triad organization in skeletal muscle and is involved in the pathophysiology of centronuclear myopathy. EMBO Molecular Medicine, 6, 1455–1475.

14. Dos Reis, R., Kornobis, E., Pereira, A., Tores, F., Carrasco, J., Gautier, C., Jahannault-Talignani, C., Nitschké, P., Muchardt, C., Schlosser, A., et al. (2022) Complex regulation of Gephyrin splicing is a determinant of inhibitory postsynaptic diversity. Nat Commun, 13, 3507.

15. Tian, B., Yang, J. and Brasier, A.R. (2012) Two-step cross-linking for analysis of protein-chromatin interactions. Methods Mol Biol, 809, 105–120.

16. Dos Reis, R., Kornobis, E., Pereira, A., Tores, F., Carrasco, J., Gautier, C., Jahannault-Talignani, C., Nitschké, P., Muchardt, C., Schlosser, A., et al. (2022) Complex regulation of Gephyrin splicing is a determinant of inhibitory postsynaptic diversity. Nat Commun, 13, 3507.

17. Allemand, E. and Ango, F. (2022) Analysis of Splicing Regulation by Third-Generation Sequencing. Methods Mol Biol, 2537, 81–95.

18. Igolkina, A.A., Zinkevich, A., Karandasheva, K.O., Popov, A.A., Selifanova, M.V., Nikolaeva, D., Tkachev, V., Penzar, D., Nikitin, D.M. and Buzdin, A. (2019) H3K4me3, H3K9ac, H3K27ac, H3K27me3 and H3K9me3 Histone Tags Suggest Distinct Regulatory Evolution of Open and Condensed Chromatin Landmarks. Cells, 8, 1034.

19. Wang, W., Carey, M. and Gralla, J.D. (1992) Polymerase II Promoter Activation: Closed Complex Formation and ATP-Driven Start Site Opening. Science, 255, 450–453.

20. Pęziński, M., Daszczuk, P., Pradhan, B.S., Lochmüller, H. and Prószyński, T.J. (2020) An improved method for culturing myotubes on laminins for the robust clustering of postsynaptic machinery. Sci Rep, 10, 4524.

21. Vilmont, V., Cadot, B., Ouanounou, G. and Gomes, E.R. (2016) A system for studying mechanisms of neuromuscular junction development and maintenance. Development, 143, 2464–2477.

22. Moransard, M., Borges, L.S., Willmann, R., Marangi, P.A., Brenner, H.R., Ferns, M.J. and Fuhrer, C. (2003) Agrin regulates rapsyn interaction with surface acetylcholine receptors, and this underlies cytoskeletal anchoring and clustering. J Biol Chem, 278, 7350–7359.

23. Oury, J., Liu, Y., Töpf, A., Todorovic, S., Hoedt, E., Preethish-Kumar, V., Neubert, T.A., Lin, W., Lochmüller, H. and Burden, S.J. (2019) MACF1 links Rapsyn to microtubule- and actin-binding proteins to maintain neuromuscular synapses. J Cell Biol, 218, 1686–1705.

24. Tintignac, L.A., Brenner, H.-R. and Rüegg, M.A. (2015) Mechanisms Regulating Neuromuscular Junction Development and Function and Causes of Muscle Wasting. Physiol Rev, 95, 809–852.

25. Tang, H., Veldman, M.B. and Goldman, D. (2006) Characterization of a muscle-specific enhancer in human MuSK promoter reveals the essential role of myogenin in controlling activity-dependent gene regulation. J Biol Chem, 281, 3943–3953.

26. Tang, H. and Goldman, D. (2006) Activity-dependent gene regulation in skeletal muscle is mediated by a histone deacetylase (HDAC)-Dach2-myogenin signal transduction cascade. Proc Natl Acad Sci U S A, 103, 16977–16982.

27. Segatto, M., Fittipaldi, R., Pin, F., Sartori, R., Dae Ko, K., Zare, H., Fenizia, C., Zanchettin, G., Pierobon, E.S., Hatakeyama, S., et al. (2017) Epigenetic targeting of bromodomain protein BRD4 counteracts cancer cachexia and prolongs survival. Nat Commun, 8, 1707.

28. Yang, N., Das, D., Shankar, S.R., Goy, P.-A., Guccione, E. and Taneja, R. (2022) An interplay between BRD4 and G9a regulates skeletal myogenesis. Front. Cell Dev. Biol., 10.

29. Haberland, M., Arnold, M.A., McAnally, J., Phan, D., Kim, Y. and Olson, E.N. (2007) Regulation of HDAC9 gene expression by MEF2 establishes a negative-feedback loop in the transcriptional circuitry of muscle differentiation. Mol. Cell. Biol., 27, 518–525.

30. Tosic, M., Allen, A., Willmann, D., Lepper, C., Kim, J., Duteil, D. and Schüle, R. (2018) Lsd1 regulates skeletal muscle regeneration and directs the fate of satellite cells. Nat Commun, 9, 366.

31. Mazhar, S. and Herbst, R. (2012) The formation of complex acetylcholine receptor clusters requires MuSK kinase activity and structural information from the MuSK extracellular domain. Molecular and Cellular Neuroscience, 49, 475–486.

32. Haddix, S.G., Lee, Y. il, Kornegay, J.N. and Thompson, W.J. (2018) Cycles of myofiber degeneration and regeneration lead to remodeling of the neuromuscular junction in two mammalian models of Duchenne muscular dystrophy. PLoS One, 13, e0205926.

33. Li, Y., Lee, Y. il and Thompson, W.J. (2011) Changes in aging mouse neuromuscular junctions are explained by degeneration and regeneration of muscle fiber segments at the synapse. J Neurosci, 31, 14910–14919.

34. Slater, C.R. (2020) ‘Fragmentation’ of NMJs: a sign of degeneration or regeneration? A long journey with many junctions. Neuroscience, 439, 28–40.

35. Valdez, G., Tapia, J.C., Lichtman, J.W., Fox, M.A. and Sanes, J.R. (2012) Shared Resistance to Aging and ALS in Neuromuscular Junctions of Specific Muscles. PLOS ONE, 7, e34640.

36. Traoré, M., Noviello, C., Vergnol, A., Gentil, C., Halliez, M., Saillard, L., Gelin, M., Forand, A., Lemaitre, M., Guesmia, Z., et al. (2024) GDF5 as a rejuvenating treatment for age-related neuromuscular failure. Brain, 147, 3834–3848.

